# The HAQ STING mutation interacts with COPA causing allosteric structural rearrangements in COPA that may protect against COPA disease

**DOI:** 10.1101/2025.03.07.642091

**Authors:** Steven Lehrer

## Abstract

**Background:** COPA disease is a rare autoinflammatory disorder caused by mutations in the COPA gene, leading to immune dysregulation and inflammatory pathology. The HAQ STING allele has been identified as a protective factor against disease manifestation. Understanding the molecular interaction between HAQ STING and COPA is critical for uncovering potential therapeutic strategies.

**Methods:** This study utilized AlphaFold2 Multimer to predict the structural interaction between COPA, STING, and HAQ STING. PyMOL was used for molecular visualization and analysis of conformational differences in protein-protein interactions. Structural alignment and binding pocket analysis were conducted to assess the potential impact of HAQ STING on COPA function.

**Results:** AlphaFold2 Multimer analysis revealed that HAQ STING causes a 90-degree rotational shift in its binding orientation to COPA compared to STING, inducing significant conformational rearrangements. The interaction alters COPA’s structural stability, suggesting an allosteric regulatory mechanism. A potential binding channel for a small therapeutic molecule was identified at the interface of STING and COPA. If the channel in the COPA STING interface meets criteria for depth, stability, and functional significance, it could be a drug-binding pocket. A therapeutic small molecule docked in a binding pocket at the interface of two interacting proteins can disrupt the protein-protein complex. This approach, known as protein-protein interaction (PPI) inhibition, is a well-established strategy in drug discovery. Fosfomycin, a phosphate-containing molecule, docked in the channel at the interface of COPA and STING.

**Conclusion:** This study provides novel structural insights into the protective role of HAQ STING in COPA disease. The conformational shift induced by HAQ STING may modulate immune signaling, preventing disease manifestation. A potential binding channel for a small therapeutic molecule was identified at the interface of STING and COPA; fosfomycin docked in the channel. These findings highlight potential therapeutic avenues, including gene therapy, small-molecule inhibitors, and STING pathway modulation. Further experimental validation is needed to translate these structural insights into clinical applications.

---

COPA disease (COPA syndrome) is a rare autoinflammatory disorder caused by mutations in the COPA gene, which encodes a protein involved in intracellular protein transport. COPA stands for Coatomer Protein Complex Subunit Alpha. It is a component of the coat protein complex I (COPI), which is involved in intracellular protein trafficking, particularly in transporting proteins from the Golgi apparatus back to the endoplasmic reticulum (ER).

COPA disease is classified as an autosomal dominant disorder and is associated with immune dysregulation. COPA Disease has multiple manifestations: 1. Lung Involvement, interstitial lung disease (ILD), particularly pulmonary hemorrhage or fibrosis. Symptoms include chronic cough and shortness of breath. 2. Kidney disease, glomerulonephritis, often with proteinuria and hematuria, often leading to kidney dysfunction. 3. Joint Involvement, arthritis (rheumatoid-like), often severe, presenting with synovitis and joint pain. 4. Immune System Dysregulation and autoantibodies, often resembling systemic autoimmune diseases like lupus or rheumatoid arthritis, as well as hyperactivation of immune pathways e.g., STING (Stimulator of Interferon Genes) pathway activation. COPA disease results from mutations in the COPA gene and defective retrograde transport of proteins between the Golgi and the endoplasmic reticulum (ER), resulting in mislocalization of immune signaling proteins, leading to chronic inflammation [1].

Simchoni et al investigated the role of the HAQ STING allele in the clinical penetrance of COPA disease. The three mutations forming the HAQ allele are R71H: Arginine (R) at position 71 → Histidine (H); G230A: Glycine (G) at position 230 → Alanine (A); and R293Q: Arginine (R) at position 293 → Glutamine (Q). Simchoni et al found that the HAQ allele co-segregated with clinical nonpenetrance in 35 individuals with COPA mutations. Experimentally, they found that HAQ STING acted dominantly to dampen COPA-dependent STING signaling. Expressing HAQ STING in patient cells abrogated the molecular phenotype of COPA disease [2]. In sum, carriers of HAQ STING were at significantly reduced risk of COPA disease.

In the current study, AlphaFold2 Multimer version 3 was used to assess the interaction between the COPA, STING, and HAQ STING proteins. Molecular Docking, a method which predicts the preferred orientation of one molecule to a second when a ligand and a target (COPA) are bound to each other to form a stable complex, was used to identify a ligand that might be a treatment for COPA disease.

## Methods

AlphaFold2 Multimer (AF2M) is an extension of the AlphaFold2 protein structure prediction tool, specifically designed to predict the structures of protein complexes (multimers). AF2M predicts how multiple protein chains assemble and interact to form a complex. This function is crucial for understanding biological processes, as many proteins function in groups [3].

Pymol, a molecular visualization tool designed for rendering and analyzing 3D molecular structures, was used for structural analysis. Pymol is extensively used in structural biology, drug discovery, and bioinformatics for visualizing proteins, nucleic acids, and small molecules. The *super* command in Pymol was used to perform structural alignment between two molecular structures by optimizing the root-mean-square deviation (RMSD) between their atomic coordinates. This command is an enhanced version of *align*, incorporating additional dynamic programming to improve alignment quality.

Molecular docking was done with AutoDock Vina Extended on the SAMSON platform (OneAngstrom, Grenoble, France). SAMSON is an interface for molecular design that has an open architecture and applicability for drug design [4]. AutoDock Vina Extended achieves approximately 2 orders of magnitude acceleration compared with the molecular docking software AutoDock 4 while also significantly improving the accuracy of the binding mode predictions. Further speed is achieved from parallelism by using multithreading on multicore machines. AutoDock Vina Extended automatically calculates the grid maps and clusters the results in a way transparent to the user [5].

## Results

Figure 1 illustrates the COPA protein domains. The WD40 domain is crucial for COPA function in vesicle trafficking. Mutations causing COPA disease are within amino acids 230-243. The WD-associated region in COPA is likely involved in stabilizing interactions within the coatomer complex, binding cargo proteins, and ensuring efficient vesicle trafficking. Both STING and HAQ STING complex with COPA at the WD40 domain.

**Figure 1.**
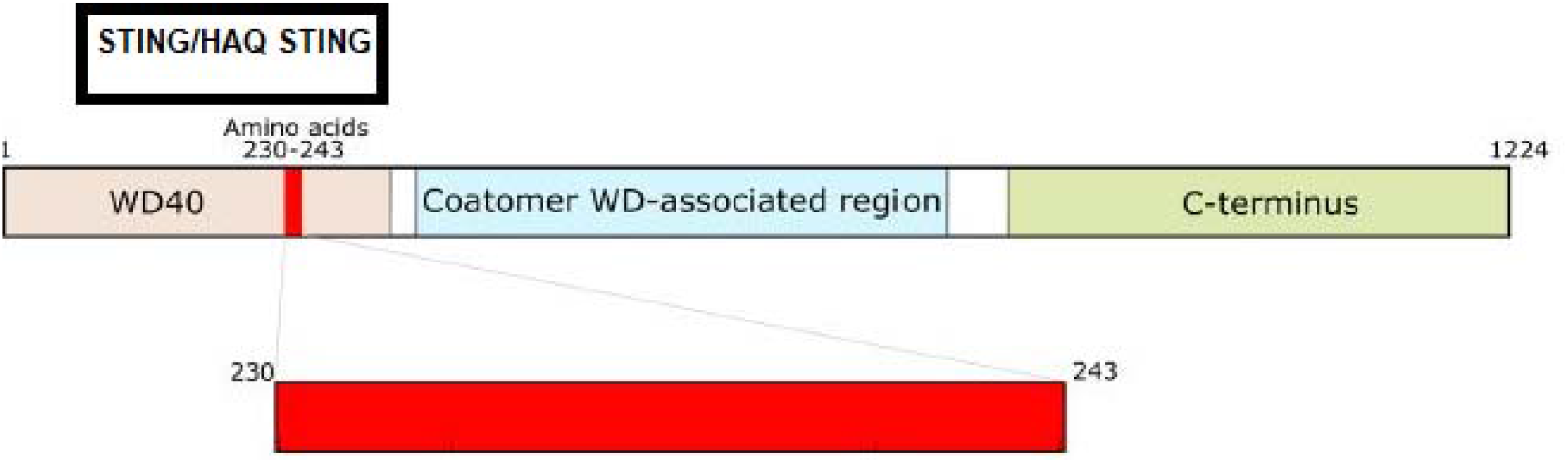
COPA protein domains. The WD40 domain is crucial for COPA function in vesicle trafficking and is composed of 285 amino acids. Mutations causing COPA disease are within amino acids 230-243. The Coatomer WD-associated region in COPA is likely involved in stabilizing interactions within the coatomer complex, binding cargo proteins, and ensuring efficient vesicle trafficking. The coatomer complex itself is a protein complex involved in intracellular vesicle transport. Both STING and HAQ STING complex with the WD40 domain. The black rectangle indicates the position of STING/HAQ STING in the protein-protein complex. Both STING and HAQ STING bind to COPA at the CDN binding domain of STING.

Figure 2 shows AF2M (Multiple Sequence Alignment) sequence coverage plot. The plot provides a visual representation of how well the input sequence (COPA/HAQ STING) aligns with other related sequences across the entire sequence. Except for positions 500 and 1220, a significant portion of the protein complex has moderate sequence identity.

**Figure 2.**
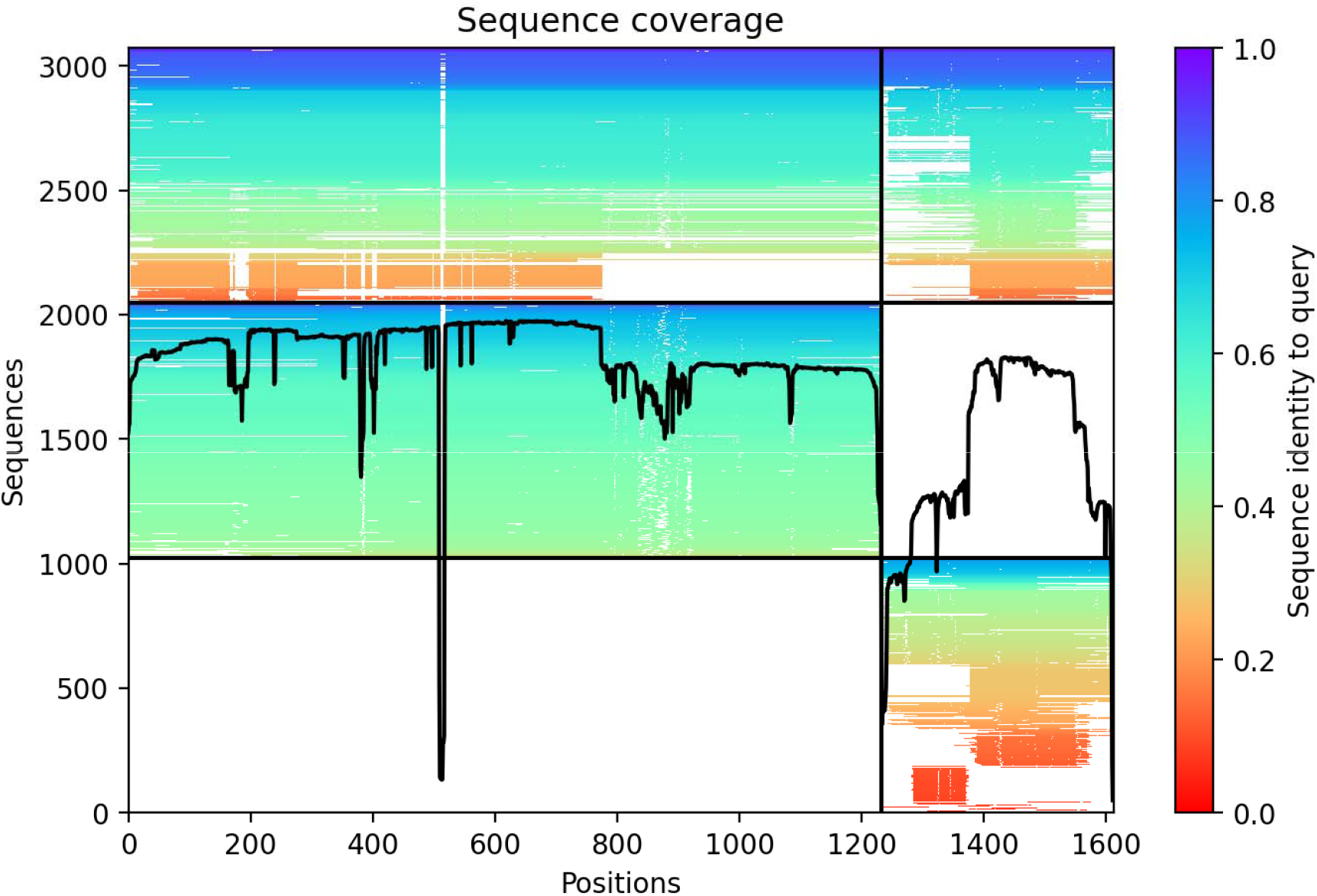
AF2M sequence coverage plot. The plot provides a visual representation of how well the input sequence aligns with other related sequences across the entire sequence. Except at positions 500 and 1220 a significant portion of the protein has moderate sequence identity.

An AF2M run with 3 recycles produced a pLDDT score (Predicted Local Distance Difference Test) ∼75-76 across all recycles, suggesting moderate confidence in the structural prediction.

pTM (Predicted TM-score, Template Modeling Score) is a global similarity measure AlphaFold2 MSA uses to compare protein structures. pTM assesses how well a predicted structure aligns with a reference e.g., an experimentally determined structure or a native fold. A native fold refers to the biologically active, three-dimensional structure of a protein under physiological conditions. It is the most stable and functional conformation of a protein, as determined by its amino acid sequence. pTM for COPA/HAQ STING initially was 0.427 and fluctuated around 0.40 after recycles. TM-score < 0.5 typically indicates low reliability in the global fold. The global fold of a protein refers to the overall three-dimensional arrangement of its entire structure, including how secondary structure elements (α-helices, β-sheets, loops) are positioned relative to each other to form a stable, functional conformation. The overall COPA/HAQ STING predicted that protein-protein interface may be unstable or variable.

Figure 3 shows predicted Aligned Error (PAE) matrices for five ranked models generated by AF2M. PAE plots help evaluate structural confidence and inter-domain flexibility. PAE measures the uncertainty in the relative positioning of different regions in the protein. All five models show similar trends, with clear high confidence folding (blue) for individual chains (COPA, HAQ STING). However, inter-domain interactions have high uncertainty (red regions off-diagonal). The individual chains are likely correctly folded but the relative orientation of subunits is not well-defined.

**Figure 3.**
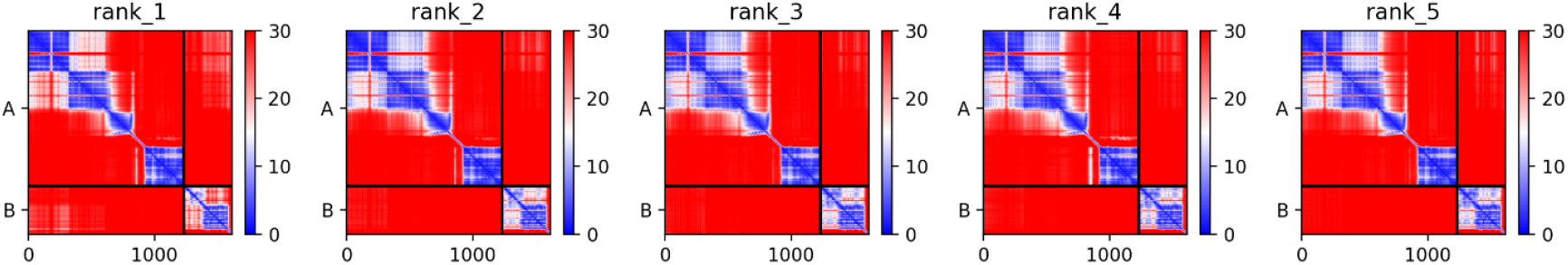
Predicted Aligned Error (PAE) matrices for five ranked models generated by AF2M. PAE plots help evaluate structural confidence and inter-domain flexibility. PAE measures the uncertainty in the relative positioning of different regions in the protein. The color scale: Blue (Low PAE = 0-10) → High confidence in spatial positioning. Red (High PAE > 20-30) → Low confidence, meaning the relative positioning of these regions is uncertain. The X and Y axes correspond to residues in the protein. The black boxes separate different chains/domains. Diagonal blue regions indicate high-confidence intra-domain folding and suggest that individual domains (A and B) are well-defined. Red off-diagonal regions indicate low-confidence relative positioning between different domains and suggest high flexibility or alternative conformations between the domains. All five models show similar trends, with clear high-confidence folding (blue) for individual chains (COPA, HAQ STING). However, inter-domain interactions have high uncertainty (red regions off-diagonal). Model 1 (rank_1) appears slightly more structured compared to lower-ranked models. The individual chains are likely correctly folded but the relative orientation of subunits is not well-defined.

Figure 4 shows predicted IDDT (intrinsic Distance Difference Test) per residue position for the top five ranked AF2M models. The pIDDT score provides a per-residue confidence measure in terms of local structure reliability. Most residues have high confidence (pIDDT > 70-90), indicating reliable local structure. Several regions show significant dips (pIDDT < 50), suggesting flexible loops, intrinsically disordered regions, poorly constrained or weakly interacting domains with possible alternative conformations.

**Figure 4.**
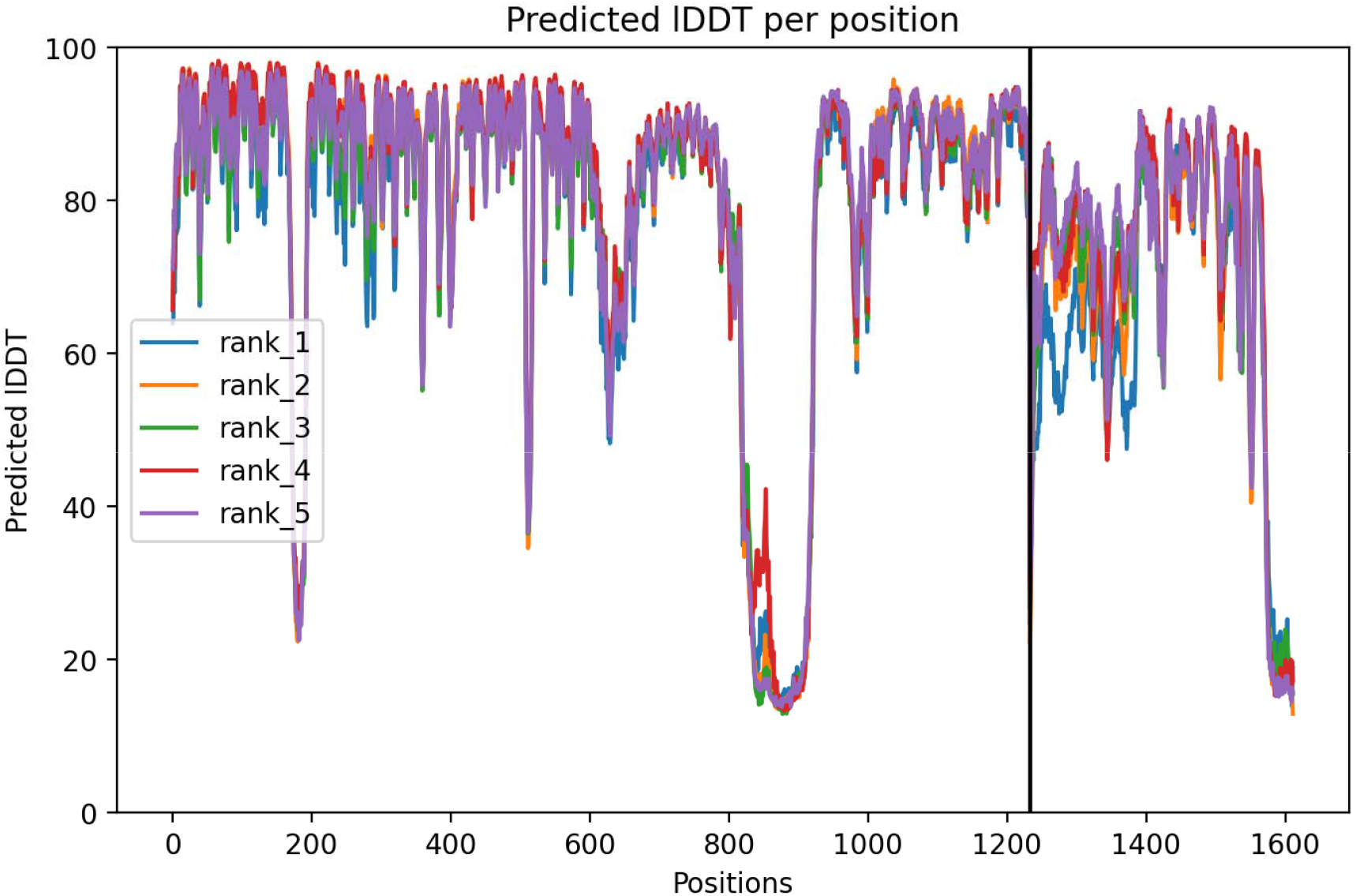
Predicted IDDT (intrinsic Distance Difference Test) per residue position for the top five ranked AF2M models. The pIDDT score provides a per-residue confidence measure in terms of local structure reliability. Most residues have high confidence (pIDDT > 70-90), indicating reliable local structure. Several regions show significant dips (pIDDT < 50), suggesting flexible loops, intrinsically disordered regions, poorly constrained or weakly interacting domains with possible alternative conformations. Black vertical line (Position 1220) marks a domain boundary or inter-protein interaction site. Fluctuations around this region suggest possible structural uncertainty in interactions.

Figure 5A shows protein-protein interaction of COPA with HAQ STING. Both STING and HAQ STING bind to COPA at the Cyclic Dinucleotide Binding Domain (CDN) of STING, around alanine 161.

**Figure 5.**
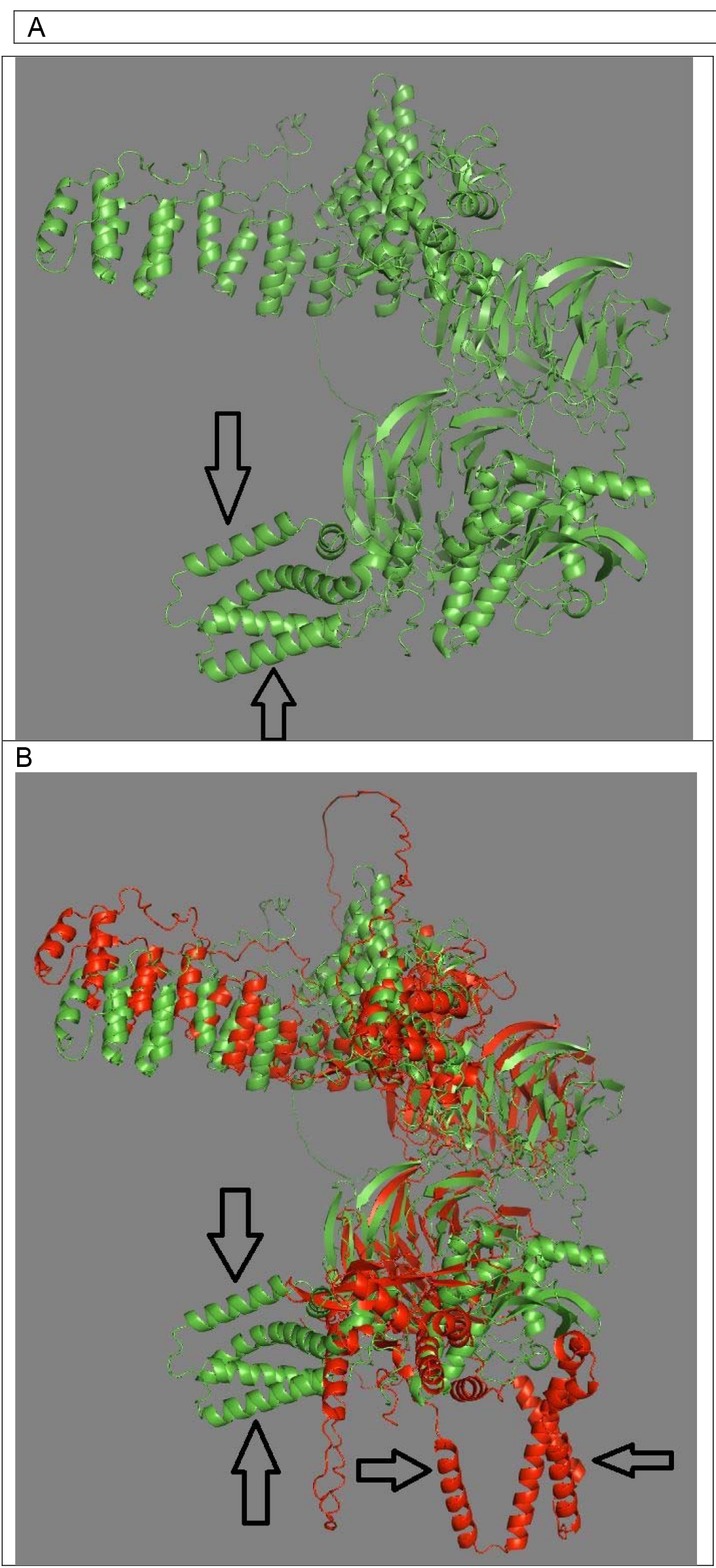
A. Protein-protein interaction of COPA with HAQ STING (up and down arrows, HAQ Sting). 5B. Protein-protein interactions of COPA with STING and HAQ STING. Two COPA protein molecules are superimposed. One COPA molecule (red) has interacted with STING (horizontal arrows). The other COPA molecule (green) has interacted with HAQ STING (up and down arrows). Note that the conformation of COPA has changed, depending on whether the interaction is with STING or HAQ STING. Pymol calculated RMSD (Root Mean Square Deviation) of 9.264 Å when aligning the two protein-protein interaction structures using the *super* command, confirming the significant conformational difference between the two complexes related to the HAQ STING rotation and altered conformation of COPA.

Figure 5B shows protein-protein interactions of COPA with STING and HAQ STING. The two COPA protein molecules are superimposed. One COPA molecule has interacted with STING. The other COPA molecule has interacted with HAQ STING. The HAQ STING interaction with COPA has rotated HAQ STING 90 degrees from the STING interaction with COPA. The conformation of COPA changes, depending on whether the interaction is with STING or HAQ STING. Both STING and HAQ STING bind to COPA at the CDN binding domain of STING.

Pymol calculated RMSD (Root Mean Square Deviation) of 9.264 Å when aligning the two protein-protein interaction structures using the *super* command, confirming the significant allosteric conformational difference between the STING COPA and HAQ STING COPA complexes related to the HAQ STING rotation and altered conformation of COPA.

Figure 6 shows WD40 domain of COPA. Two large potential binding channels are present based on their spatial distribution relative to the protein’s center. Largest potential channel distance is ∼50.17 Å from the center of mass, second largest channel distance is ∼49.52 Å from the center of mass. These distances (∼50 Å) suggest significant cavities, which could potentially accommodate therapeutic small molecule docking.

**Figure 6.**
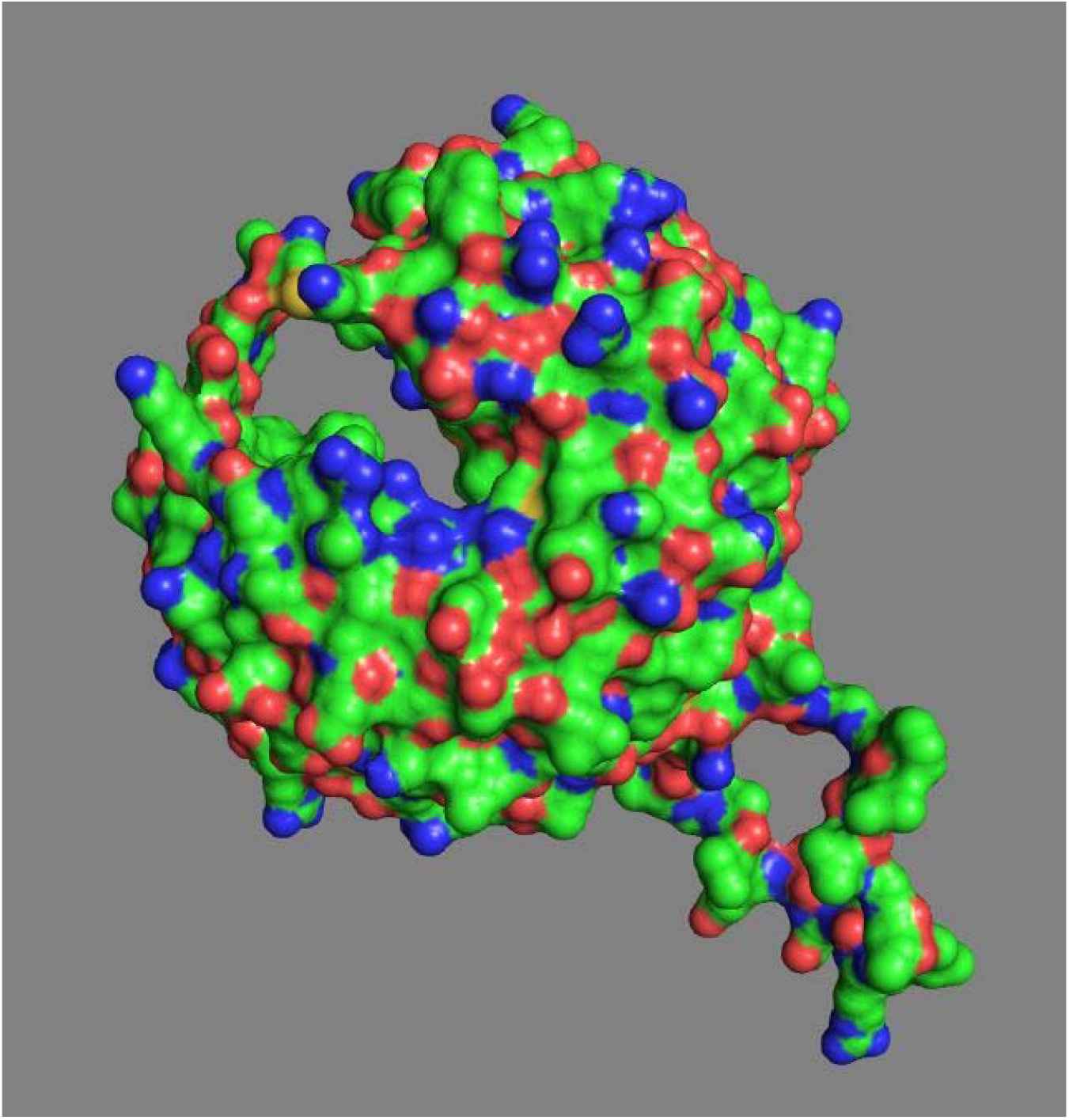
WD40 domain of COPA. Two large potential binding channels are present based on their spatial distribution relative to the protein’s center. Largest Potential Channel (upper left) Distance: ∼50.17 Å from the center of mass, Second Largest Channel (lower right) Distance: ∼49.52 Å from the center of mass. These distances (∼50 Å) suggest significant cavities, which could potentially accommodate therapeutic small molecule docking. The three colors are hydrophobicity coloring: Green → Moderately hydrophilic or neutral residues, Blue → Highly hydrophilic residues (polar, charged), Red → Hydrophobic residues (nonpolar, like Leucine, Isoleucine, Valine).

Figure 7A shows COPA STING interface, COPA above, STING below. Note channel at center of interface, which might serve as a protein-protein interface (PPI) binding pocket. The channel could be a binding site for a small therapeutic molecule. Figure 7B shows the COPA HAQ STING interface. Note that the channel at center of interface is absent.

**Figure 7.**
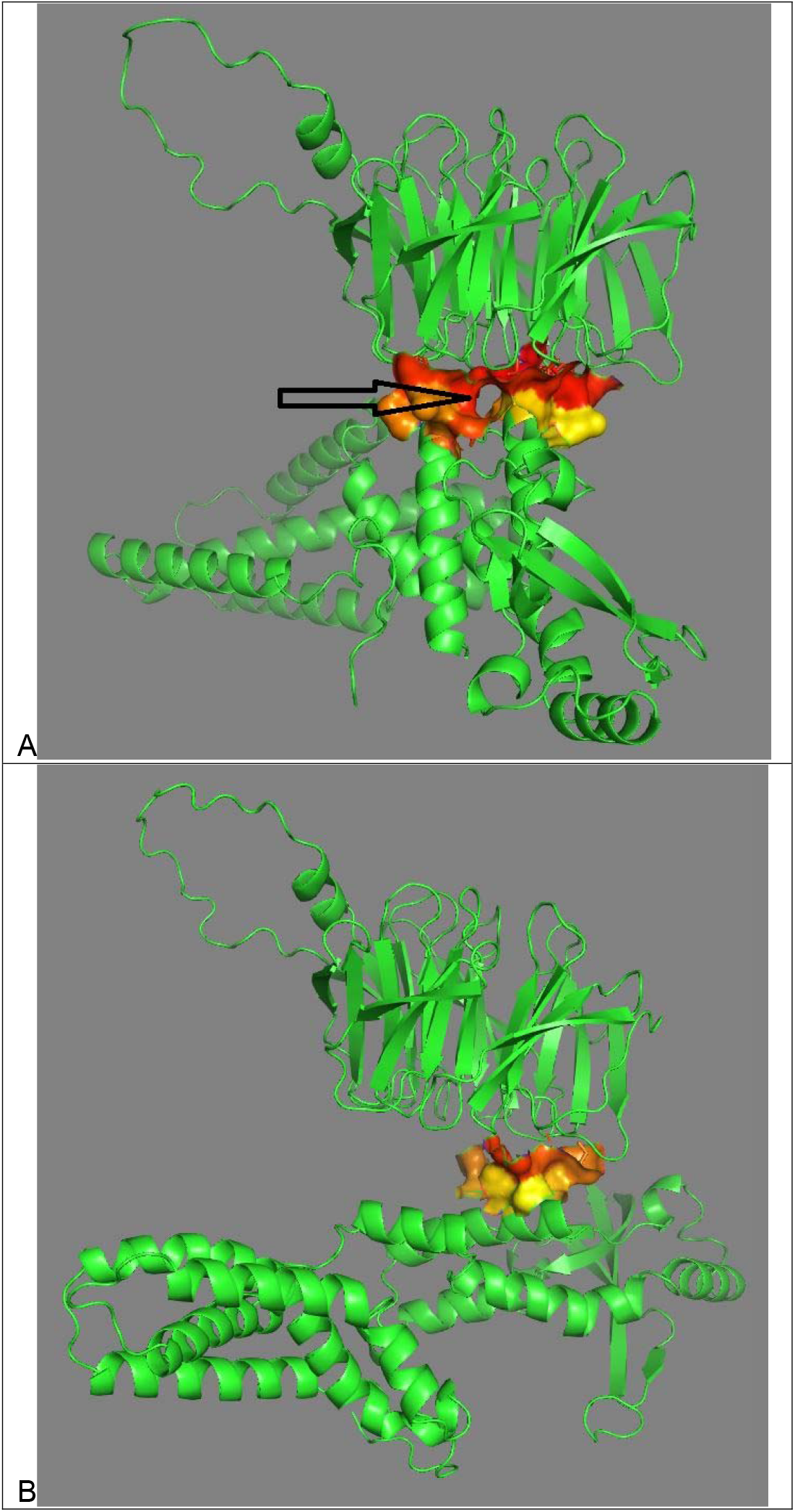
Structural Comparison of COPA-STING and COPA-HAQ STING Interfaces. (A) COPA-STING interface: 3D structural rendering of the interaction between the COPA WD40 domain (top) and STING (bottom), highlighting a central binding channel (indicated by arrow) at the interface. This channel may serve as a potential protein-protein interaction (PPI) binding pocket suitable for small-molecule drug targeting. (B) COPA-HAQ STING interface: Structural rendering of the same region showing the interaction between COPA and the HAQ variant of STING. Notably, the previously observed central channel at the COPA-STING interface is absent here, indicating a significant conformational rearrangement induced by HAQ STING binding. The difference suggests an allosteric shift in protein orientation that may contribute to HAQ STING’s protective effect.

Figure 8 shows a Fosfomycin molecule docked in the channel at the interface of COPA and STING. Table 1 lists the molecular docking parameters. Mode 1, the best and most reliable binding pose, is illustrated in Figure 8. These data support the structural plausibility of fosfomycin or related phosphate-containing molecules occupying the identified PPI pocket.

**Table 1.** Docking results from AutoDock Vina Extended showing ten predicted binding modes of fosfomycin within the COPA-STING interface. Mode 1 exhibits the strongest binding affinity (−4.12 kcal/mol) and zero RMSD, indicating a stable and well-defined docking pose. Affinity (kcal/mol) reflects binding strength (more negative = stronger binding). Ki (μmol) represents the predicted inhibition constant, with lower values indicating higher potency. RMSD (lb and ub) denote lower and upper bounds of positional deviation from reference poses. These data support the structural plausibility of fosfomycin or related phosphate-containing molecules occupying the identified PPI pocket.

| mode | affinity (kcal/mol) | ki (μmol) | RMSD (lb Å) | RMSD (ub Å) |
| --- | --- | --- | --- | --- |
| 1 | -4.119699645 | 955.5193 | 0 | 0 |
| 2 | -4.114005744 | 964.7463 | 9.122865296 | 10.1468622 |
| 3 | -4.103604042 | 981.833 | 9.617445436 | 10.5307809 |
| 4 | -4.098236494 | 990.7682 | 9.080182153 | 10.1035447 |
| 5 | -4.068585633 | 1041.613 | 9.644921736 | 10.4563062 |
| 6 | -3.994190534 | 1180.968 | 8.063276036 | 8.8669117 |
| 7 | -3.889636599 | 1408.889 | 8.898417432 | 9.91143621 |
| 8 | -3.773483376 | 1714.027 | 9.153883982 | 10.0160061 |
| 9 | -3.743448626 | 1803.156 | 8.767985011 | 9.72495238 |
| 10 | -3.73895452 | 1816.885 | 9.62578458 | 10.5981295 |

**Figure 8.**
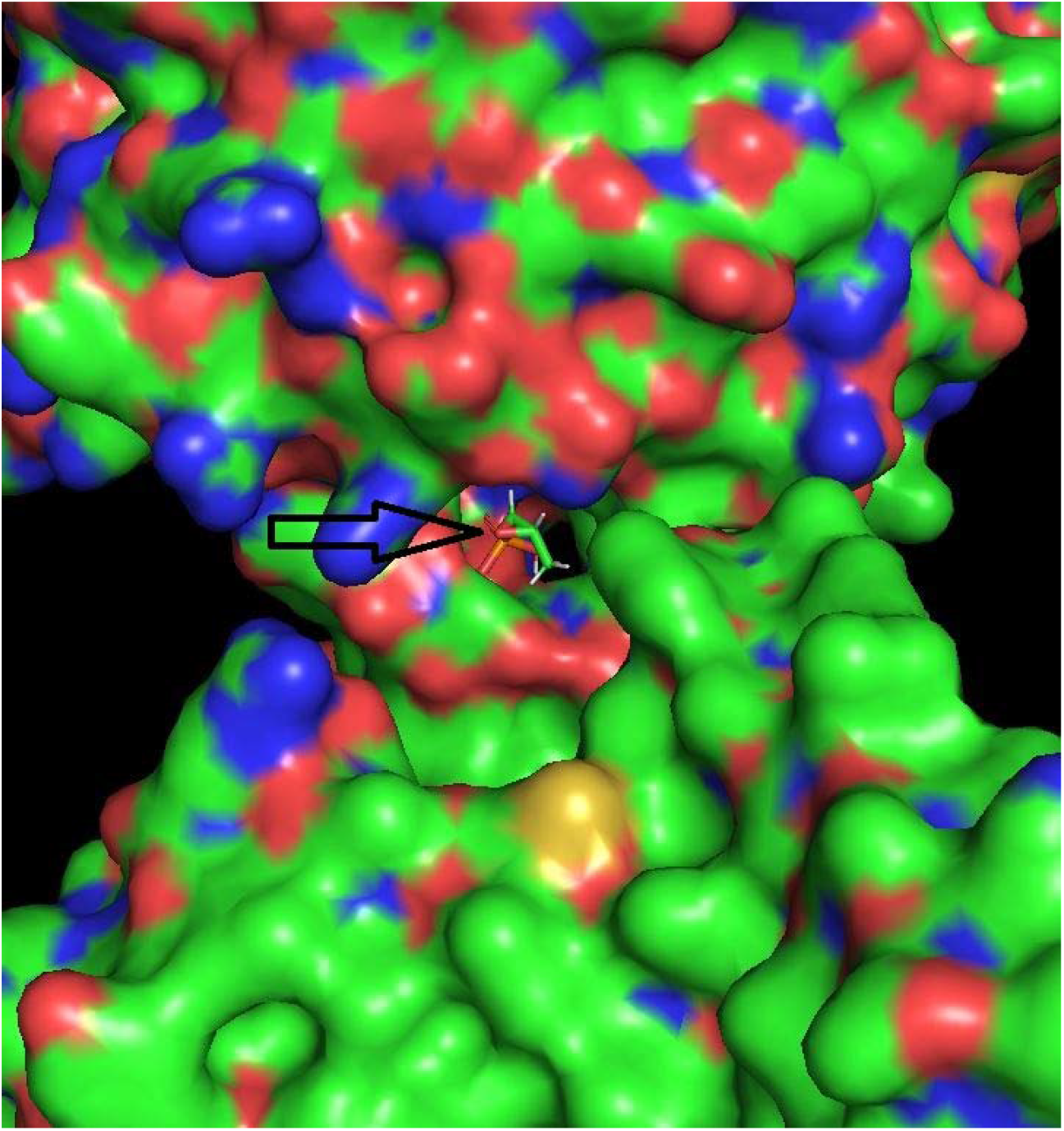
Docking of Fosfomycin into the COPA-STING Binding Channel. Molecular docking visualization shows a fosfomycin molecule (highlighted with an arrow) positioned within the channel at the interface of COPA (above) and STING (below). This suggests the feasibility of targeting this pocket with small-molecule inhibitors. The docking pose represents Mode 1, the most energetically favorable configuration based on AutoDock Vina Extended results. The presence of positively charged arginine residues within the channel may facilitate electrostatic interactions with the phosphate group of fosfomycin.

## Discussion

In an AlphaFold2 protein-protein interaction (PPI) prediction, if a mutation causes one protein to rotate 90 degrees relative to its orientation with another protein, a significant change in the binding interface and interaction dynamics has taken place. In this case, induced conformational change has occurred. The HAQ STING mutation has caused structural rearrangements in COPA and STING proteins, altering their preferred binding orientation. The rearrangements may suggest allosteric effects, where a distant mutation changes the overall shape and binding properties.

Because STING signaling is implicated in COPA disease pathology, STING is a prime therapeutic target. The Cyclic Dinucleotide Binding Domain (CDN) (Residues: 151–339) of both STING and HAQ STING complexes with COPA around STING residue alanine 160. The CDN domain binds cyclic dinucleotides (CDNs), such as cyclic GMP-AMP (cGAMP), leading to the activation of downstream signaling pathways.

STING inhibitors target the cyclic dinucleotide binding pocket [6]. The pocket is located primarily within the Cyclic Dinucleotide Binding Domain. A small molecule drug in this pocket that produced STING loss of function could result in reduced inflammation, which may be beneficial in autoinflammatory disorders such as COPA disease. Both STING and HAQ STING bind to COPA at the CDN binding domain of STING.

Small-molecule STING inhibitors are in clinical development and may help reduce inflammation [7]. However, these inhibitors could increase infection risk, raising concerns about their safety in patients.

Other current therapeutic approaches to COPA disease are multifaceted. Immunosuppressive agents such as corticosteroids reduce inflammation during disease exacerbations. Disease-Modifying Antirheumatic Drugs (DMARDs) like methotrexate and azathioprine serve as maintenance therapies to control symptoms. Rituximab, a monoclonal antibody targeting CD20-positive B cells, is used in some cases to manage severe symptoms [1]. Janus Kinase (JAK) small molecule inhibitors Baricitinib and Ruxolitinib have shown promise by modulating the overactive interferon signaling pathway [2].

Future therapeutic approaches could involve genetic modification of HAQ STING. Since HAQ STING is a protective genetic variant that prevents disease manifestation in COPA mutation carriers, a potential gene therapy approach would be to introduce HAQ STING into patients’ cells, a universal therapeutic strategy.

STING Trafficking Modulation could represent another therapy [8]. STING trafficking refers to the movement of the STING protein within a cell, particularly between cellular compartments like the endoplasmic reticulum (ER), Golgi apparatus, and lysosomes. This trafficking process is crucial for regulating immune responses, especially in conditions involving chronic inflammation, autoimmune diseases, and infections. The Simchoni et al study highlights the importance of STING’s localization in the Golgi apparatus as a disease mechanism [2].

Therapies that alter STING trafficking might be explored as a treatment strategy. The channel in the COPA STING interface (Figure 7B) can be considered a potential binding pocket for small-molecule drug development if the channel meets certain criteria for depth, stability, and functional significance. Identifying and characterizing such pockets is a crucial step in structure-based drug design [9]. A therapeutic small molecule docked in a binding pocket at the interface of two interacting proteins can disrupt the protein-protein complex. This approach, known as protein-protein interaction (PPI) inhibition, is a well-established strategy in drug discovery [10, 11].

The COPA Sting interface channel contains arginine (ARG), which can interact with phosphate-containing molecules such as fosfomycin in biological systems. Arginine (ARG) is positively charged, has a guanidinium group, and interacts strongly with negatively charged phosphate groups (e.g., in DNA, ATP, phosphorylated proteins). Therefore, the docking of fosfomycin within the COPA Sting interface channel is not surprising. Experimental screening is needed to confirm if fosfomycin or a similar molecule can effectively modulate Copa disease.

Given the role of COPA protein’s WD40 repeat domain in HAQ STING protein-protein interactions and the COPA STING interface potential binding pocket described above, designing small molecules that can modulate the protein-protein interaction presents an additional potential therapeutic avenue. Such targeted therapies could significantly improve patient outcomes.

The study has weaknesses, especially lack of experimental validation. The findings are based entirely on in silico predictions. Without biochemical or cellular validation (e.g., co-immunoprecipitation, cytokine profiling, or reporter assays), the conclusions remain speculative. While AlphaFold2 is cutting-edge, the low pTM (∼0.40) and high PAE in inter-domain regions suggest instability or variability in the COPA-HAQ STING interface. The large RMSD (9.26 Å) should be interpreted with caution. Fosfomycin was chosen as a docked molecule based on charge complementarity rather than biological relevance to COPA or STING. Broader virtual screening of more disease-relevant compounds would strengthen the therapeutic argument.

Structural findings are extrapolated to propose STING signaling suppression without direct evidence of immune modulation. No data is available on downstream pathway activity (e.g., interferon response).

Nevertheless, the study emphasizes the importance of structural bioinformatics tools such as AlphaFold2 in uncovering disease mechanisms and identifying therapeutic targets. The ability to model protein-protein interactions at high resolution provides insights into molecular pathophysiology, guiding rational drug design and precision medicine approaches.

## Conclusion

Structural analysis using AlphaFold2 Multimer highlights the critical role of HAQ STING in modifying COPA conformation and mitigating COPA disease pathology. The observed rotational shift in HAQ STING binding to COPA suggests an allosteric regulatory mechanism that may suppress pathogenic immune activation. The channel in the COPA STING interface can be considered a potential binding pocket for small-molecule therapeutic drug development if the channel meets certain criteria for depth, stability, and functional significance. Fosfomycin docks within this channel. These findings pave the way for novel therapeutic approaches, including small-molecule inhibitors, gene therapy, and STING pathway modulation. Further experimental validation and preclinical studies are essential to translate these structural insights into clinical applications for COPA disease treatment. The integration of computational modeling with biochemical and genetic studies will be crucial in advancing our understanding of COPA disease and developing targeted therapies to improve patient outcomes.

